# Chondroitin Sulfate Flourishes Gut Sulfatase-Secreting Bacteria To Damage Mucus Layers, Leak Bacterial Debris, And Trigger Inflammatory Lesions In Mice

**DOI:** 10.1101/145714

**Authors:** Tao Liao, Yan-Ping Chen, Li-Li Tan, Chang-Qing Li, Qi Wang, Shui-Qing Huang, Xin-An Huang, Qin Xu, Qing-Ping Zeng

## Abstract

**Background:** An interaction of the food types with the gut microbiota changes is deeply implicated in human health and disease. To verify whether animal-based diets would lead to gut dysbiosis, systemic inflammation and inflammatory pathogenesis, we fed mice with chondroitin sulfate (CS), a sulfate-containing *O*-glycan naturally occurring in livestock and poultry products, and monitored the dynamic changes of microbial flores, inflammatory signatures, and pathogenic hallmarks.

**Results:** A metagenomic gut microbiota analysis revealed the overgrowth of sulfatase-secreting bacteria and sulfate-reducing bacteria in the gastrointestinal tracts of mice upon daily CS feeding. Sulfatase-secreting bacteria compromise gut integrity through prompting mucin degradation and mucus lesions, which were evident from the upregulation of secretary leukocyte protease inhibitor (SLPI) and mucin 1/4 (MUC-1/4). A synchronous elevation of lipopolysaccharide (LPS) and tumor necrosis factor α (TNF-α) levels in the serum as well as cerebral, hepatic, cardiac and muscular tissues suggests bacterial endotoxinemia, chronic low-grade inflammation and mitochondrial dysfunction, eventually leading to the onset of global inflammatory pathogenesis towards arthritis, dementia, tumor, and fatty liver.

**Conclusions:** CS triggers the early-phase and multi-systemic pathogenesis like arthritis, dementia, tumor, and fatty liver by enhancing gut opportunistic infection and evoking low-grade inflammation in mice. A plausible reason for the inconsistency of CS in treatment of osteoarthritis (OA) was also discussed.

## Background

Human and animal gastrointestinal tracts are colonized by commensal microbes, mainly including bacteria and fungi, with differential and food-dependent gut microbiota compositions among individuals [1]. An animal-based high fat/high protein food intake increases the abundance of *Bacteroides*, whereas a plant-based high fiber food intake increases the richness of *Prevotella* [2,3]. While a high fat/high sugar diet used in the developed "Western" nations increases the bile tolerant *Bacteroides, Alistips* and *Bilophila*, a high fiber diet eaten by the people in developing countries increases the polysaccharide degradable *Firmicutes*, such as *Roseburia, Eubacterium, rectale* and *Ruminococcus bromii* [4]. Children from a rural African village show a unique abundance of cellulose-degradable *Prevotella* and xylan hydrolyzable *Xylanibacter*, completely lacking in children from an urban European city [5].

Mounting evidence supports an alteration of the gut microbiota in patients with stroke [6], obesity [7,8] and type 2 diabetes [9,10]. A recent work has validated that the altered intestinal microbiome is related to the dysfunctional motor phenotype in patients with Parkinson’s disease [11]. Although an implication of a specific gut microbiota in human health and disease remains ambiguous, progresses regarding food-induced disorders have been achieved from the most recent findings. For example, metabolism of L-carnitine, a trimethylamine abundant in red meat, by the intestinal microbiota was found to accelerate atherosclerosis by conversion of L-carnitine to trimethylamine-N-oxide in mice [12]. Chondroitin sulfate (CS), a sulfate-containing ingredient in animal products including red meat, was shown to promote the overgrowth of sulfate-reducing bacteria (SRB) in aid of sulfatase-secreting bacteria (SSB), during which sulfatase degrades CS as an electric recipient for sulfate reduction [13]. The gut microbiota flourished by heme, another red meat component, was also noticed to facilitate an abnormal epithelial proliferation via reducing the S-S bond for mucolysis and opening the mucus barrier in the colon [14].

Given that an interaction of CS with SSB allows the bacterial endotoxin lipopolysaccharide (LPS) trespassing the gut barrier and entering the blood stream, it might be anticipated that CS should trigger inflammatory responses and aggravate inflammatory conditions such as inflammatory bowel disease (IBD) and osteoarthritis (OA). However, a randomized and controlled clinical trial for the treatment of cannie IBD indicated that oral CS and probiotics (resistant starch, β-glucans and mannaoligosaccharide) show improvement in selected serum biomarkers, hence assuming a reduction in disease activity [15]. Indeed, CS in combination with glucosamine or glucosamine sulfate has long been used for the dietary intervention of OA [16]. Nevertheless, it was also warned that CS should not be used to treat the symptomatic OA because of without relief [17]. Recently, it was observed that a combined therapy of CS with glucosamine sulfate or glucosamine hydrochloride exhibits no improvement of joint damage in a rabbit OA model [18], and CS plus glucosamine sulfate shows no superiority over placebo in a randomized, double-blind and placebo-controlled clinical trial in OA patients [19]. Nevertheless, a question regarding why CS is unlikely always effective on OA remains unanswered.

Considering CS impacting on the gut microbiota homeostasis described above [13], we proposed here that CS might exert distinctive effects on OA in a gut microbiota-dependent manner. In other words, CS should improve OA if gut SSB is absent, otherwise CS might aggravate OA. This is because SSB, in addition to degrade CS, can also degrade mucin-containing mucus linings, by which LPS should be leaked out from the gut. Actually, some CS-degrading bacteria, including *thetaiotaomicron* J1 and 82, *B. ovitus* E3 and *Clostridium hathewayi* R4, were isolated and characterized from healthy persons [20]. Additionally, it was proved that the gut microbiota is more vulnerable to CS in female mice than male mice [21]. We also assumed that the beneficial effect of CS on OA might be an essential consequence of LPS depletion by the neutralizing anti-LPS antibodies, after which a serum test would give a negative result of the inflammatory indicators.

It could be predicted that CS-induced opportunistic infection and enhanced mucus leakage should lead to LPS entering into the blood circulation, but anti-LPS not neutralizing all leaked LPS. Thus, an essential outcome should be that the residual LPS triggers multi-systemic inflammation and eventually causes inflammatory disorders. As supporting evidence, intranasal LPS infusion was shown to induce neuroinflammation and Parkinson’s disease in rodents [22]. Our previous work also revealed that articulate LPS injections can induce synovitis, an early-phase presentation of rheumatoid arthritis [23]. In theory, LPS-elicited chronic inflammation could also lead to other inflammatory consequences, including benign tumor and even malignant cancer albeit occasionally. By identifying the gut microbiota spectra in CS-fed mice and monitoring the expression profiles of selective biomarkers involving dementia, arthritis, tumor and fatty liver, we established an association of gut dysbiosis with the multi-loci inflammatory pathogenesis. These results should shed light on elucidation of a mechanistic link of the disturbed gut microbiota to some dietary factors as a major etiological originator initiating chronic inflammation and metabolic diseases.

## Methods

### Animals and treatment

The female Kunming mice (nine-month old, 40-45 g) that belong to an out-bred population from SWISS mice were provided by The Experimental Animal Centre of Guangzhou University of Chinese Medicine in China (Certificate No. 44005800001448). All mice were housed on a 12-h light and 12-h dark cycle at 25°C with *ad libitum* chow and free water drinking. After three-week quarantine, mice were randomly divided into a control group fed only with *ad libitum* chow (AL mice), a model group intragastrically administered with 250 μl CS (100 mg/ml, FocusChem, Shandong, China) daily for two months in addition to *Ad libitum* feeding (CS mice), a model group intragastrically administered with 250 μl CS (FocusChem, Shandong, China) and 104 *Bacillus cereus* (Huankai Microbial Sci & Tech Co Ltd., Guangdong, China) daily for two months in addition to *Ad libitum* feeding (CS-BC mice), and a model group peritoneally injected with 0.25 mg/kg LPS (Sigma Aldrich) on every two days for two weeks in addition to *Ad libitum* feeding (LPS mice). Three to six mice were included within each group. All experiments were approved by The Animal Care Welfare Committee of Guangzhou University of Chinese Medicine (No. SPF-2015009). The experimental protocols complies with the requirements of animal ethics issued in the Guide for the Care and Use of Laboratory Animals of the National Institute of Health (NIH), USA.

### Gut microbiota metagenomic analysis

The gut microbiota in a mouse fecal sample was identified by the high-throughput 16S VX sequencing-based classification procedure. The sequencing (sample preparation, DNA extraction and detection, amplicon purification, library construction and online sequencing) and data analysis (paired end-reads assembly and quality control, operational taxonomic units cluster and species annotation, alpha diversity and beta diversity) were conducted by Novogene, Beijing, China.

### Polymerase chain reaction (PCR) microarray and quantitative PCR (qPCR)

The RT^2^ Profiler^™^ PCR Array Mouse Fatty Liver (PAMM-157Z) was purchased from SABioscience Qiagen, Hilden, Germany. The experiment was performed by Kangchen Biotechnology Co, Ltd, Shanghai, China. RNA isolation, purity determination, electrophoresis monitoring, reverse transcription, and quantification were performed obeying a standard protocol. The relative copy numbers = 2^−ΔΔCᵀ^, in which ΔCᵀ = Ct _target gene_ - Ct _reference gene_, ΔΔCᵀ = ΔC _treatment sample_ - ΔC _control sample_. Each pair of primers listed in **Additional file 1** were designed and applied under the following amplification condition: 95°C, 30s; 95°C, 3s, 64°C, 34s, 45 cycles.

### Western blotting (WB) and enzyme-linked immunosorbent assay (ELISA)

The reference protein glyceraldehyde-3-phosphate dehydrogenase (GAPDH), and all antigen proteins were immunoquantified by WB according to manufacture’s manuals. Antibodies against mouse mucin 1 (MUC1)/MUC4, cyclin D1 (CD1), and p21 were purchased from CapitalBio Corp, China. ELISA kits for mouse tumor necrosis factor α (TNF-α), TNF-α receptor 1 (TNFR1), toll-like receptor 4 (TRL4), chemokine (C-C motif) ligand 2 (CCL2), amyloid β peptide (Aβ) and tyrosine hydroxylase (TH) were from Beijing Chenglin Biotech Co., China. ELISA kits for mouse TNF-α, LPS, presenilin 1 (PS-1), rheumatoid factor (RF) was from Shanghai Jinma Experiment Equip Co., Ltd., China.

### Histochemical analysis by haematoxylin-eosin (HE) staining

Fix the dissected tissue by immersion in a 10% formalin solution for 4 to 8 hours at room temperature. Mount in OCT embedding compound, and freeze at -20 to -80°C. Cut 5-15 μm thick tissue sections using a cryostat. Thaw-mount the sections onto gelatin-coated histological slides. Slides are pre-coated with gelatin to enhance adhesion of the tissue. Dry the slides for 30 min on a slide warmer at 37°C. Sections were deparaffinized by xylene, re-hydrated by gradient alcohol, and washed in distilled water. After haematoxylin staining, wash in running tap water, differentiate in 1% acid alcohol, wash again in running tap water, blue in 1% ammonia, wash again in running tap water, and rinse in 95% alcohol. After eosin counter staining, dehydrate through gradient alcohol, clear in xylene, and mount with xylene-based mounting medium.

### Histochemical analysis by oil red O staining

For oil red O staining, cut fresh frozen tissue sections at 5-10 μm thick and mount on slides. Air dry slides for 30-60 min at room temperature and then fix in ice cold 10% formalin for 5-10 min. Air dry again for another 30-60 min or rinse immediately in 3 changes of distilled water. Let slides air dry for a few min. Place in absolute propylene glycol for 2-5 min to avoid carrying water into Oil Red O. Stain in pre-warmed 0.5% oil red O solution (0.5 g oil red O and 100 ml propylene glycol) for 8-10 min in 60 ºC oven. Differentiate in 85% propylene glycol solution for 2-5 min. Rinse in 2 changes of distilled water. Stain in haematoxylin for 30 sec. Wash thoroughly in running tap water for 3 min. Place slides in distilled water. Mount with glycerin jelly.

### Immunohistochemical analysis

The primary antibody against secretory leukocyte protease inhibitor (SLPI) was provided by Novus. Formalin-fixation, paraffin-embedment and deparaffinization were the same as HE staining procedure described above. Sections were incubated at room temperature with 3% H2O2 to block endogenous peroxidase, and then repaired in boiling citric acid. After washing in phosphate-buffered solution, sections were blocked by 2% bovine serum albumin and incubated with 1:100 diluted primary antibodies at 37°C for 1 h. After washing again, sections were incubated with biotinylated secondary antibodies at 37°C for 20 min. After washing again, sections were incubated with diaminobenzidine for 1-5 min. After rinsing with tap water, sections were counter stained by haematoxylin. After completion of dehydration, clearance and mounting, pictures were taken under the microscope (OLYMUPUS BX-51).

### Electronic microscopy and spectrophotometry

After treatment, cells were harvested and fixed in 2.5% glutaraldehyde in 0.1 M phosphate buffer for three hours at 4 °C, followed by post-fixation in 1% osmium tetroxide for one hour. Samples were dehydrated in a graded series of ethanol baths, and infiltrated and embedded in Spurr’s low-viscosity medium. Ultra-thin sections of 60 nM were cut in a Leica microtome, double-stained with uranyl acetate and lead acetate, and examined in a Hitachi 7700 transmission electron microscope at an accelerating voltage of 60 kV. The reagent kits of choline acetyltransferase (ChAT) for detection of activity and acetylcholine (ACh) for measurement of amounts were purchased from Nanjing Jiancheng Biotech Co., China. All determination procedures were according to manufacturers’ instructions.

### Statistical analysis

The software SPSS 22.0 was employed to analyze data, and the software GraphPad Prism 5.0 was employed to plot graphs. The Independent Simple Test was used to compare all groups, but the Kruskal-Wallis Test followed by Nemenyi test was used when the data distribution is skewed. The significance level (*p* value) was set at <0.05 (*), <0.01 (**), <0.001 (***) and <0.0001 (****).

## Results

### CS induces arthritis-like synovitis and alters gut microbiota compositions in an individual dependent manner

To verify whether a sulfate-containing nutrient might induce gut opportunistic infection through the overgrowth of some commensal bacteria, we fed AL mice daily with 25 mg of CS each. After four weeks, CS-fed mice show either a normal phenotype or an inflammatory phenotype like synovitis, an early-phase manifestation of arthritis among CS.RA, CS.HL and CS.MN. Compared with AL1 (Fig. 1A), CS.RA exhibits erythematous and edematous paws (Fig. 1B), whereas CS.HL and CS.MN do not show any distinguishable inflammatory characters. Upon histopathological analysis, it was only noticed that one or a few layers of synovia exist in the synovial tissue of AL1 (Fig. 1C), but multi-layer intimal hyperplasia and inflammatory synovial infiltration occur in the synovial tissue of CS.RA (Fig. 1D).

**Fig. 1.**
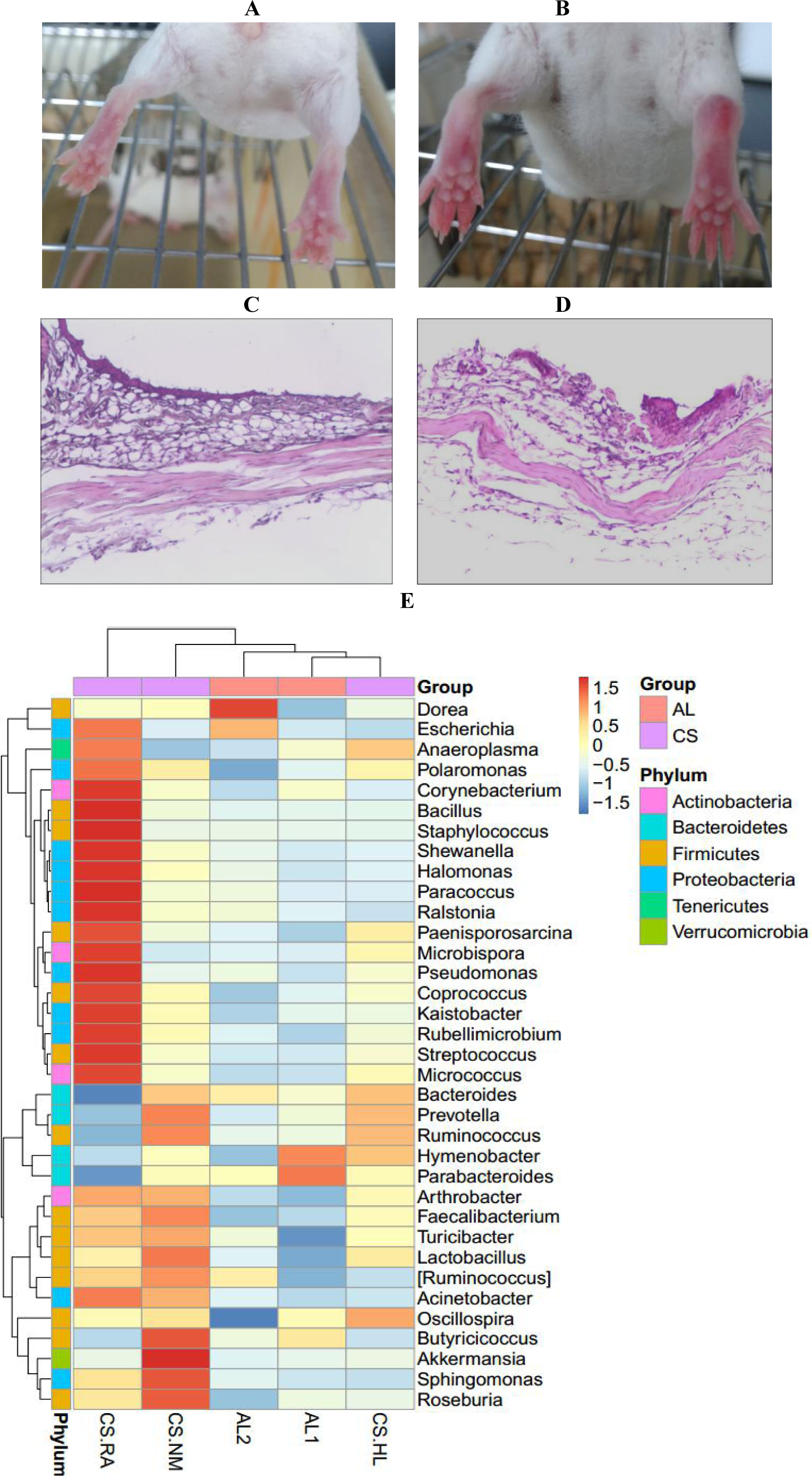
CS-induced synovial inflammation and gut microbiota fluctuation in AL and CS mice. A. AL1 without RA-like symptoms; B. CS.RA with RA-like erythematous and edematous paws; C. HE staining of synovial tissues of AL1; D. HE staining of synovial tissues of CS.RA; E. The relative abundance of bacterial phyla in the gut of AL and CS mice, in which the cold and hot colors represent Z values from -1.5 to 1.5.

To explain why CS induces different morphological changes within a same treatment group, we assumed that an inflammatory phenotype might be attributed to CS-mediated gut opportunistic infection. So we compared the gut microbiota profiles of AL and CS mice using a high-throughput 16S VX sequencing-based metagenomic analysis procedure (**Additional file 2**). As results illustrated in Fig. 1E, all five tested mice are dominantly inhabited by six commensal bacterial phyla, i.e., *Actinobacteria, Bacteroidetes, Firmicutes, Proteobacteria, Tenericutes* and *Verrucomicrobia*. However, a different composition of each phylum was evident among AL and CS mice. For example, CS.RA shows abundant *Proteobacteria* (hot/red color), including *Escherichia, Polaromonas, Shewanella, Hallomonas, Paracoccus, Ralstonia, Pheudomonas, Kaistobacter* and *Rubellimicrobium*, but rare *Bacteroidetes* (cold/blue color), such as *Bacteroides, Prevotella, Hymenobacter* and *Parabacteroides*. In contrast, CS.NM has more *Bacteroidetes* but less *Proteobacteria.* As to CS.HL, its phylum proportion is more similar with AL mice rather than CS mice. These findings indicated that not every mouse carries a completely identical gut microbiota community, and the gut microbiota composition would be changed or unchanged in an individual-dependent manner.

The abundant microbiota in the gut of CS.RA belongs to nine genera of Gram-negative *Proteobacteria* and five genera of Gram-positive *Firmicutes*, which represent two largest bacterial phyla containing SSB and SRB. As listed in Table 1, one of the SSB species *B. cereus* from *Firmicutes* is extraordinarily enriched in the gut of CS.RA (4.03%), and another SSB species *A. muciniphila* is extremely enriched in CS.NM (4.00%). In average, CS mice not only possess more SSB than AL mice (1.655% vs 0.27%), but also carry more SRB than AL mice (1.23% vs 0.86%). Therefore, SSB should have nourished SRB by degrading CS and cellular sulfate components such as mucin to provide sulfate as an electric recipient for reduction of sulfate to hydrogen sulfide (H_2_S).

**Table 1.** ClassificationofSSBandSRBinthegutmicrobiotaofALandCSmice

| | | AL1 | AL2 | $\bar{x}\pm s$ | CS.RA | CS.HL | CS.NM | $\bar{x}\pm s$ |
| --- | --- | --- | --- | --- | --- | --- | --- | --- |
|  |  | (%) | (%) | (%) | (%) | (%) | (%) | (%) |
| SSB | *A. muciniphila* | 0.40 | 0.20 | 0.30±0.14 | 0.50 | 0.50 | 4.00 | 1.67±2.02 |
|  | *B. cereus* | 0.24 | 0.24 | 0.24±0.00 | 4.03 | 0.65 | 0.24 | 1.64±2.08 |
| SRB | *Desulfovibrionaceae* | 0.97 | 0.75 | 0.86±0.16 | 1.31 | 1.08 | 1.30 | 1.23±0.13 |

These results indicated that CS-induced overgrowth of SSB and SRB might depend on a "food chain", where SSB secrete sulfatase for degradation of CS and mucin, and SRB reduce sulfate to H2S. While CS nourishes *B. cereus* and *Desulfovibrionaceae* in CS.RA, it alternatively nourishes *muciniphila* and *Desulfovibrionaceae* in CS.NM. Unfortunately, we did not know whether CS mice prior to CS feeding have the same gut microbiota with AL mice although AL1 and AL2 harbor the identical or similar SSB and SRB. Following mucin degradation, the intestinal integrity would be compromised, thereby leading to LPS leaking out from the gut and entering into the blood stream.

### CS elevates LPS and TNF-α levels and upregulates dementia and arthritis hallmarks

To ravel whether gut opportunistic infection might lead to LPS entry into the circulation, we determined the serum level of LPS in AL and CS mice. In consequences, CS.RA shows not only a higher serum level of LPS than AL1 (16.18 U/L vs 11.96 U/L), but also an elevated serum levels of the dementia marker PS-1 (144.15 pg/ml) and the arthritis marker RF (32.44 U/L) as compared to AL1 (80.82 pg/ml and 30.89 U/L), hinting that CS might have initiated a pathogenic process towards dementia and arthritis. In similar, both serum and brain TNF-α levels are synchronously elevated in CS.RA, suggesting systemic and neuroinflammatory responses (Table 2).

**Table 2.** ComparisonofLPS,PS-1,RF,andTNF-αlevels between AL and CS mice

| Tests | AL1 | AL2 | AL3 | $\bar{x}\pm s$ | CS.RA | CS.NM | CS.HL | $\bar{x}\pm s$ |
| --- | --- | --- | --- | --- | --- | --- | --- | --- |
| Serum LPS (U/L) | 11.96 | 14.94 | 16.64 | 14.51±1.93 | 16.18 | 16.09 | 13.29 | 15.19±1.64 |
| Serum PS-1 (pg/ml) | 80.82 | 220.59 | 159.44 | 153.61±70.07 | 144.15 | 116.64 | 133.67 | 131.49±13.88 |
| Serum RF | 30.89 | 37.11 | 16.80 | 28.27±10.41 | 32.44 | 27.32 | 28.69 | 29.48±2.65 |
| (U/L) |  |  |  |  |  |  |  |  |
| Serum TNF-α | 378.73 | 526.25 | 763.2 | 556.06±193.96 | 1067.18 | 407.87 | 664.67 | 713.24±332.33 |
| (ng/L) |  |  |  |  |  |  |  |  |
| Brain TNF-α | 513.50 | 611.85 | 881.40 | 668.92±190.47 | 1150.96 | 569.96 | 741.16 | 820.69±298.55 |
| (ng/L) |  |  |  |  |  |  |  |  |

To make sure if CS is exactly implicated in dementia induction by triggering neuroinflammation, we further measured the cerebral levels of Aβ, PS-1, LPS, and TNF-α in AL1 and CS.RA. Surprisingly, Aβ and TNF-α levels are elevated (Fig. 2A-2B), whereas LPS and PS-1 levels are declined in CS.RA (Fig. 2C-2D). For this result, we proposed that LPS and PS-1 levels should be converted from high levels to low ones. Indeed, we detected high serum LPS and PS-1 levels in CS mice (see Table 2). So we assumed a mechanism underlying that eradication of LPS and subsequent downregulation of PS-1 are driven by anti-LPS antibodies, during which Aβ and TNF-α are participated in activation of the immune system.

**Fig. 2.**
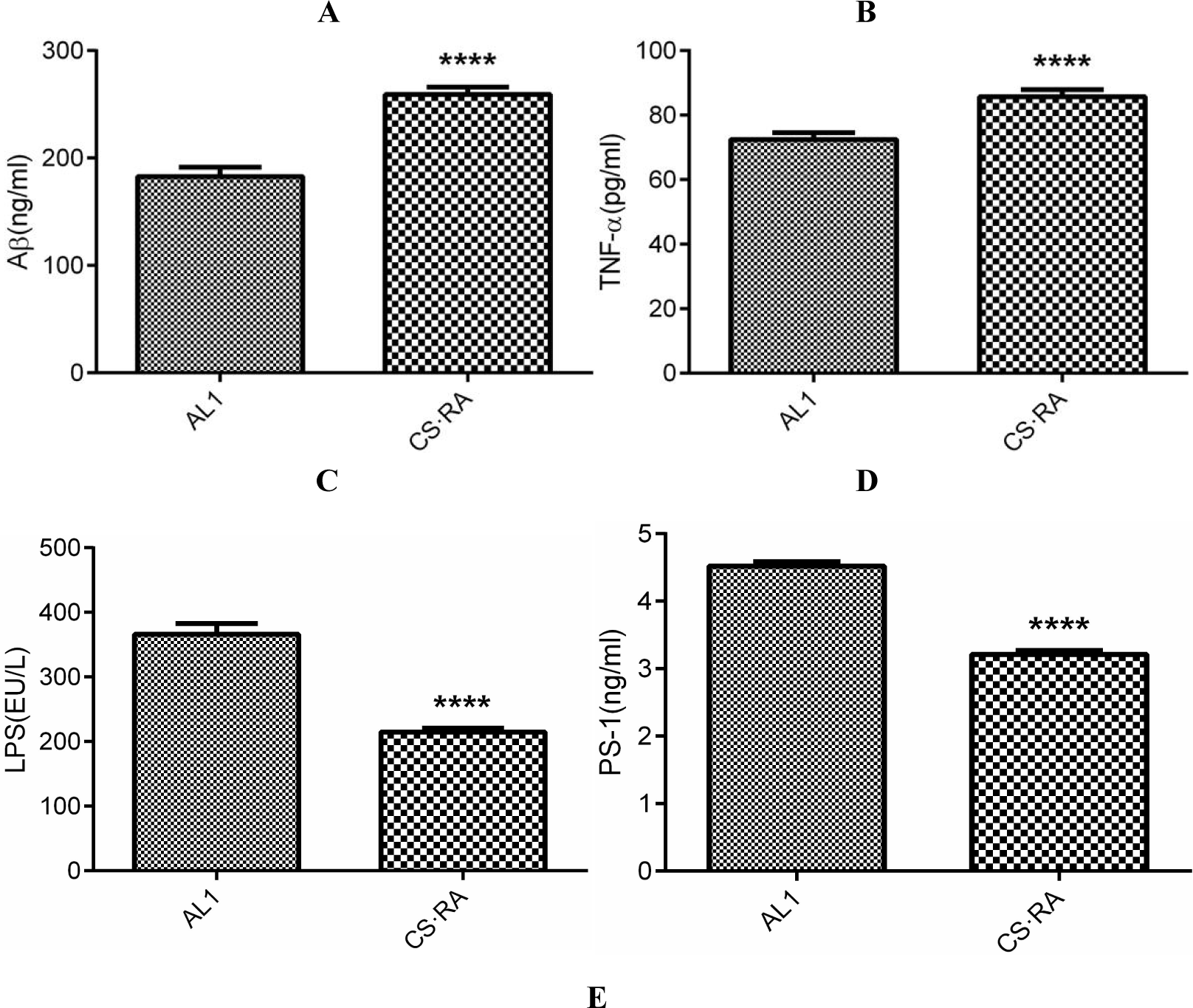
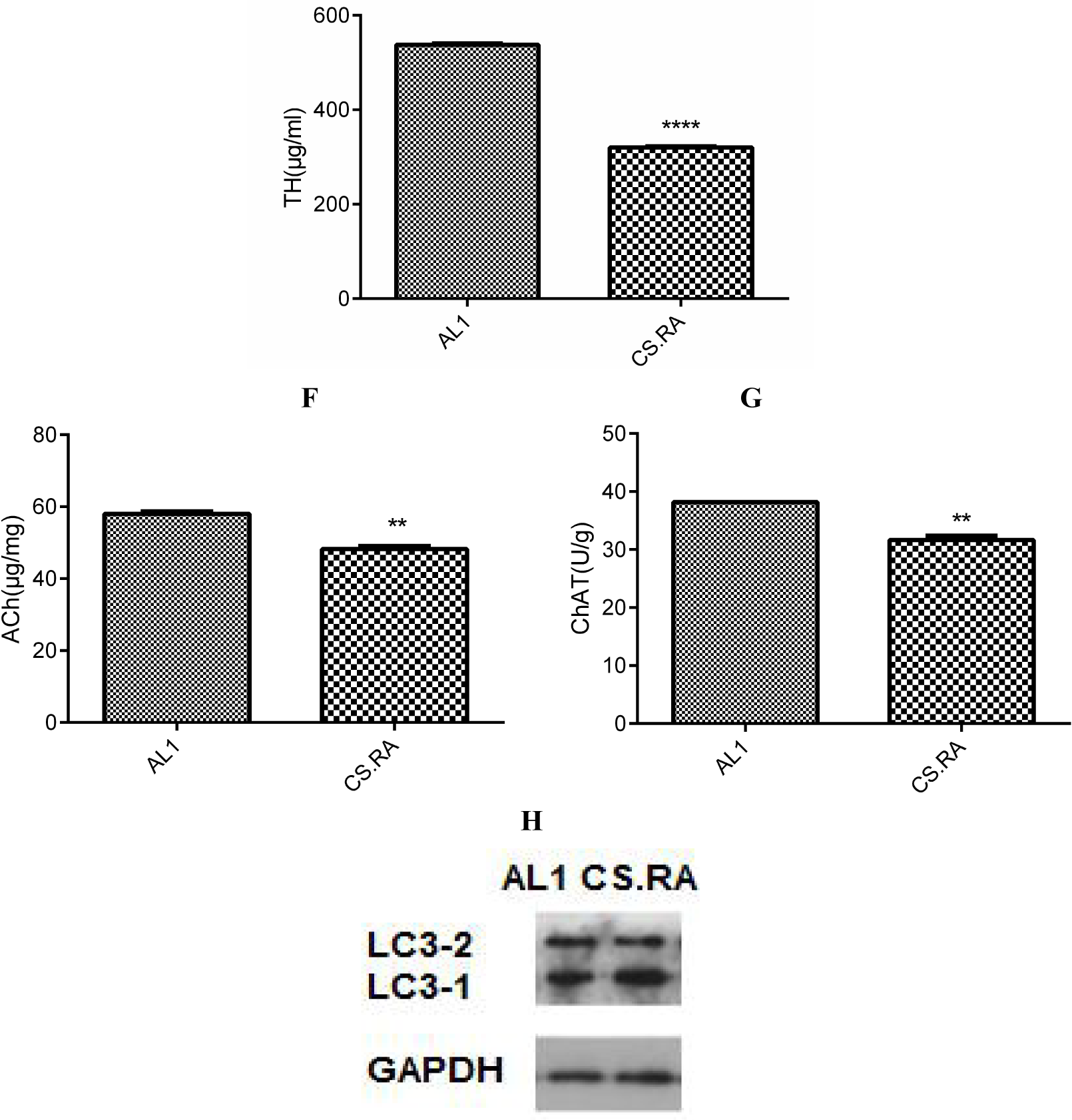
Expression of inflammatory and pathogenic markers in the cerebral tissues of AL1 and CS.RA mice. A. Aβ levels. B. TNF-α levels. C. LPS levels. D. PS-1 levels. E. TH levels; F. Ach levels; G. ChAT levels. H. LC3-2/1. **** *p*<0.0001. *** *p*<0.001.

Furthermore, we evaluated the functional changes in the brain by determining the cerebral TH, Ach and ChAT levels. Consequently, the cerebral level of TH, a rate-limiting enzyme for biosynthesis of the anti-inflammatory neurotransmitter dopamine, is lower in CS.RA than in AL1 (Fig. 2E), suggesting an attenuated anti-inflammatory ability in the brain of CS.RA. Accordingly, the cerebral levels of the cholinergic neurotransmitter Ach and its rate-limiting biosynthetic enzyme ChAT are also lower in CS.RA than in AL1 (Fig. 2F and 2G), implying a compromised neuronal function and a progressed cognitive deficit in CS.RA. It was also seen from Fig. 2H that the relative autophagy-inactivating LC3-1 level is higher than the relative autophagy-activating LC3-2 in CS.RA than AL1, highlighting that the autophagic functions are attenuated in CS.RA.

### CS induces pro/anti-inflammatory responses in a tissue specific pattern

To further elucidate whether the circulated LPS would trigger systemic inflammation, we determined the levels of pro-inflammatory and anti-inflammatory markers in different organs and tissues. As compared to AL1, CS.RA exhibits an elevated TNF-α level in hepatic and muscular tissues, but a declined TNF-α level in adipose tissues (Fig. 3A). Intriguingly, LPS levels in three tested organs are all lower in CS.RA than in AL1 (Fig. 3B). For the lower TNF-α level seen in the adipose tissue of CS.RA, we assumed that it might experience a conversion from a high level to low one.

**Fig. 3.**
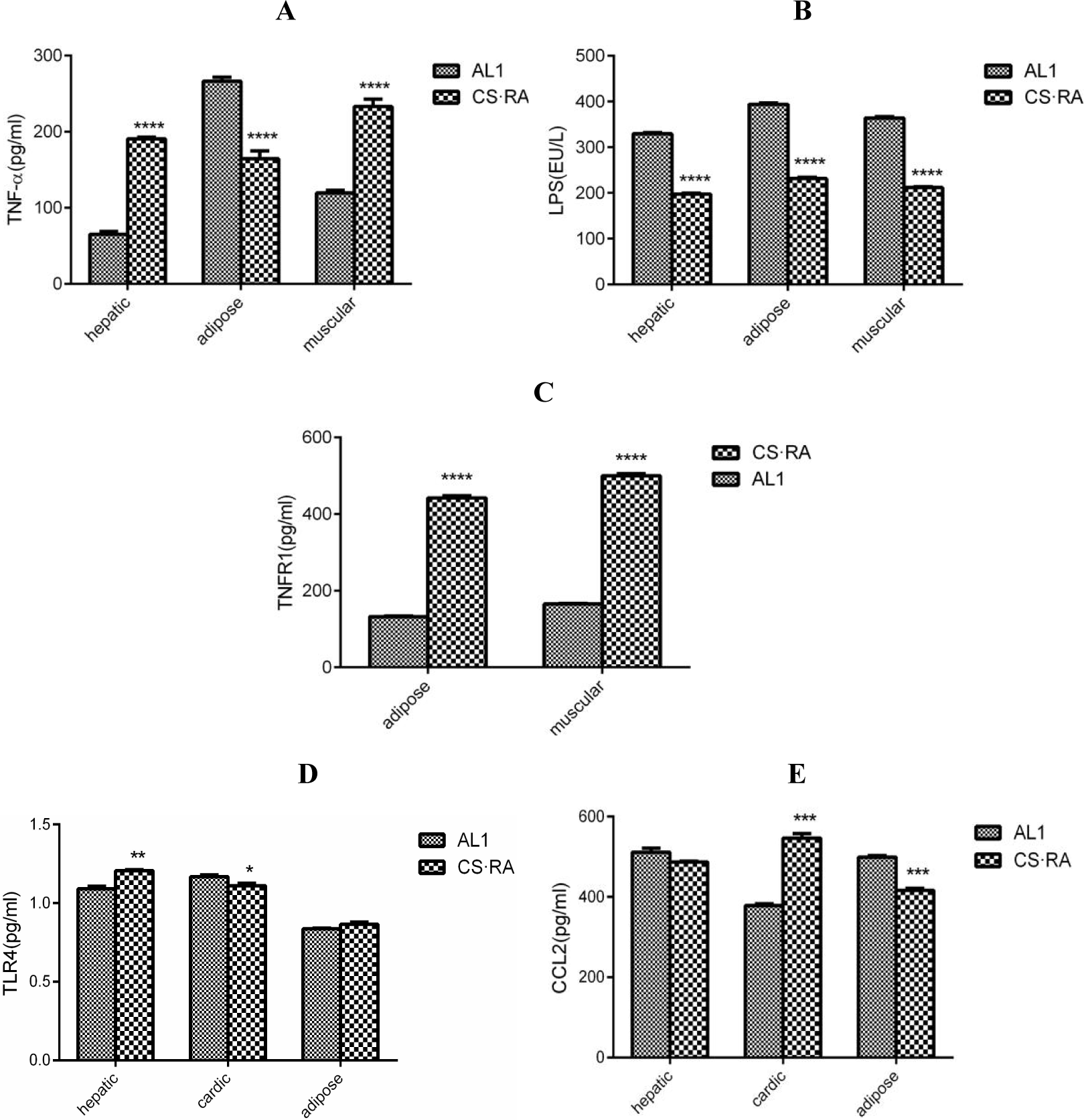
Pro/anti-inflammatory responses to CS-driven gut dysbiosis in different tissues of AL1 and CS.RA mice. A. TNF-α levels in adipose, hepatic and muscular tissues. B. LPS levels in adipose, hepatic and muscular tissues. C. TNFR levels in adipose and muscular tissues. D. TLR4 levels in hepatic, cardiac and adipose tissues. E. CCL2 levels in hepatic, cardiac and adipose tissues. **** *p*<0.0001. *** *p*<0.001. ** *p*<0.01.* *p*<0.05.

Evidence supporting this assumption is available from the comparison of TNFR1 levels among the adipose, hepatic and muscular tissues of AL1 and CS.RA mice. As illustrated in Fig. 3C, TNFR1 levels are higher in the adipose tissue, implying that LPS induces TNF-α at first, then TNF-α activates TNFR1, and TNFR1-transduced signaling finally depletes LPS in the adipose tissue. The LPS-binding receptor TLR4 is upregulated in the hepatic and cardiac tissues, but unchanged in the adipose tissue of CS.RA (Fig. 3D). At the same time, the signaling chemokine CCL2 is upregulated in the cardiac tissue, and unchanged in the hepatic tissue, or downregulated in the adipose tissue of CS.RA (Fig. 3E). These results seemed to address that adipose tissues proceed a conversion from a pro-inflammatory phase to an anti-inflammatory phase, but other tissues are maintained in a pro-inflammatory status.

### CS induces sulfatase secretion and mucin degradation accompanying with enhanced SLPI and MUC1/MUC4 expression

Although CS was shown to augment the selective inflammatory and pathogenic signatures, data listed above were available only on an individual basis. Therefore, we enlarged the scale of tested samples for further verification. To ascertain CS-induced inflammation and pathogenesis are resulted from SSB overgrowth and sulfatase overproduction, we quantified the mRNA levels of sulfatase of SSB and ATP sulfurylase of SRB in the fecal samples of AL and CS mice (Table 3). However, no difference of average values was observed between AL and CS mice.

**Table 3.** The relative copy numbers of sulfatase or ATP sulfurylase mRNAs in AL and CS mice (*n*=5)

| Fold changes of target mRNA/16S RNA |
| --- |

|  | Sulfatase | Sulfatase 1 | Sulfatase 2 | T-sulfatase | ATP sulfurylase |
| --- | --- | --- | --- | --- | --- |
| AL | 1.60±1.466 | 1.08±0.274 | 1.15±0.303 | 1.23±0.313 | 1.41±0.580 |
| CS | 1.45±0.669 | 1.12±0.278 | 1.28±0.379 | 1.56±0.837 | 1.45±0.674 |

Note: Sulfatase exists in *B. cereus*. Sulfatase 1/2 exists in *A. muciniphila*. T-sulfatase exists in *B. thetaiotaomicron*; and ATP sulfurylase exists in *D. desulfuricans*.

Because each mouse has a unique array of the gut microbiota, it should not be compared with the average levels of sulfate-degrading enzymes. Given that the lowest mRNA levels of sulfatases represent the less abundant SSB, and the highest mRNA levels of sulfatases represent the much abundant SSB, it should be compared on an individual basis. As consequences, all tested sulfatases are significantly upregulated in a CS mouse as compared with an AL mouse (Fig. 4). Interestingly, it was obviously that CS2 and CS3 show the higher sulfatase levels than other mice. While t-sulfatase mRNA of *B. thetaiotaomicron* shows the highest level in CS2, sulfatase mRNA of *B. cereus* and sulfatase 1 mRNA of *A. muciniphila* shows the highest levels in CS3. Actually, the level of sulfatase 2 mRNA of *A. muciniphila* is also higher in CS3, only slightly lower than that in CS4.

**Fig. 4.**
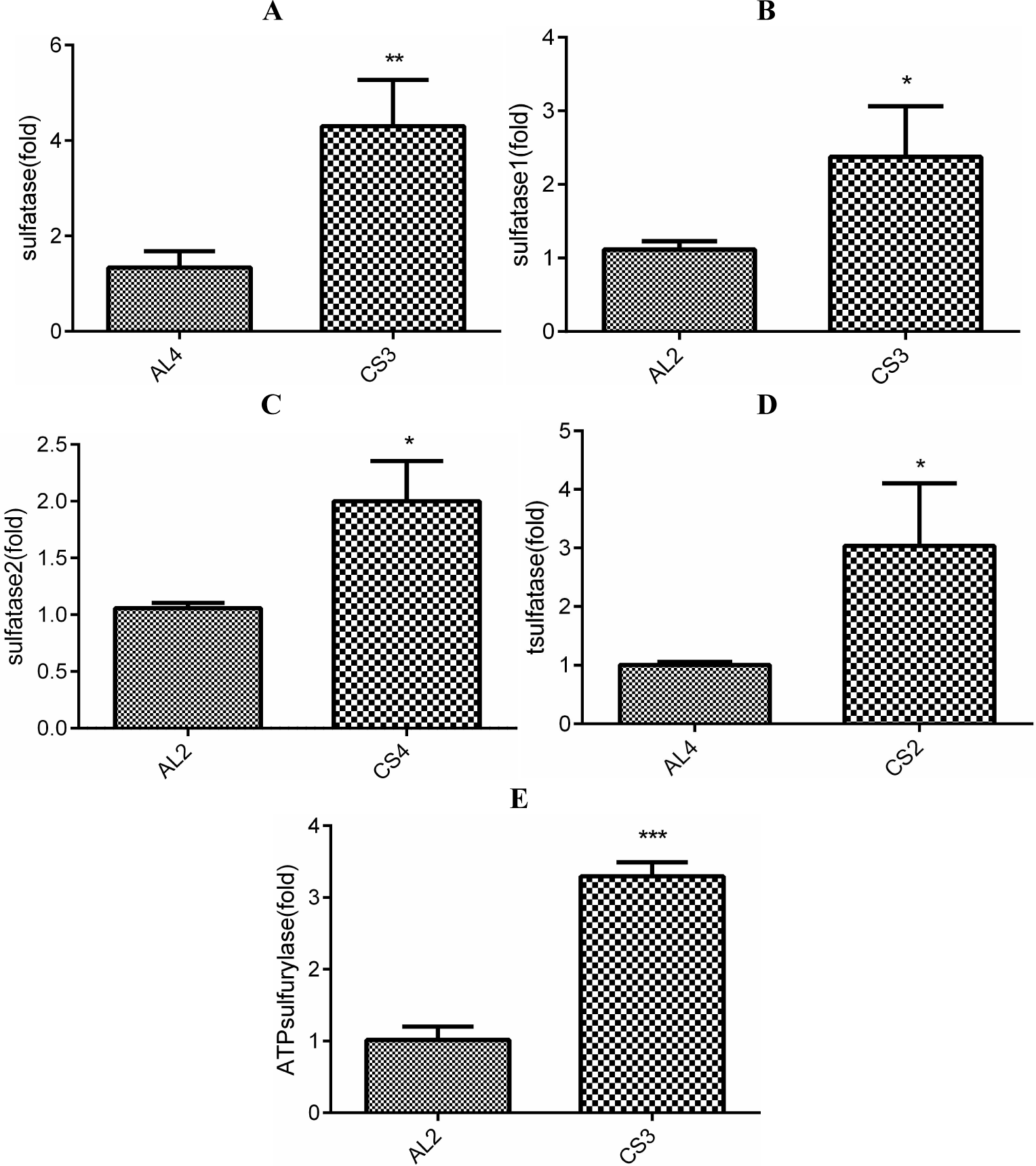
Expression of bacterial sulfatases and ATP sulfurylase in the gut microbiota of AL and CS mice. A. sulfatase mRNA levels in *B. cereus.* B. sulfatase 1 mRNA levels in *A. muciniphila.* C. sulfatase 2 mRNA levels in *A. muciniphila.* D. T-sulfatase mRNA levels in *B. thetaiotaomicron*. E. ATP sulfurylase mRNA levels in *D. desulfuricans.* *** *p*<0.001. ** *p*<0.01. * *p*<0.05.

Once mucin is degraded, SLPI should be upregulated because of mucus damage. As compared with AL2 (Fig. 5A), SLPI is remarkably upregulated in CS2 (Fig. 5B) and CS3 (Fig. 5C). Interestingly, as shown in Fig. 4, CS2 shows overexpression of sulfatase in *B. thetaiotaomicron*, and CS3 exhibits the high levels of sulfatase in *B. cereus* and sulfatase 1/2 in *A. muciniphila*, indicating an association of high sulfatase levels with enhanced SLPI expression. On the other hand, mucus damage should be also anticipated to upregulate MUC1/MUC4 for mucus repairing. Indeed, it was noticed that MUC1 is upregulated in CS3 and CS4, while MUC4 is upregulated in CS1, CS2 and CS3 (Fig. 5D). Unexpectedly, MUC1 and MUC4 are upregulated in AL3 with an unknown reason.

**Fig. 5.**
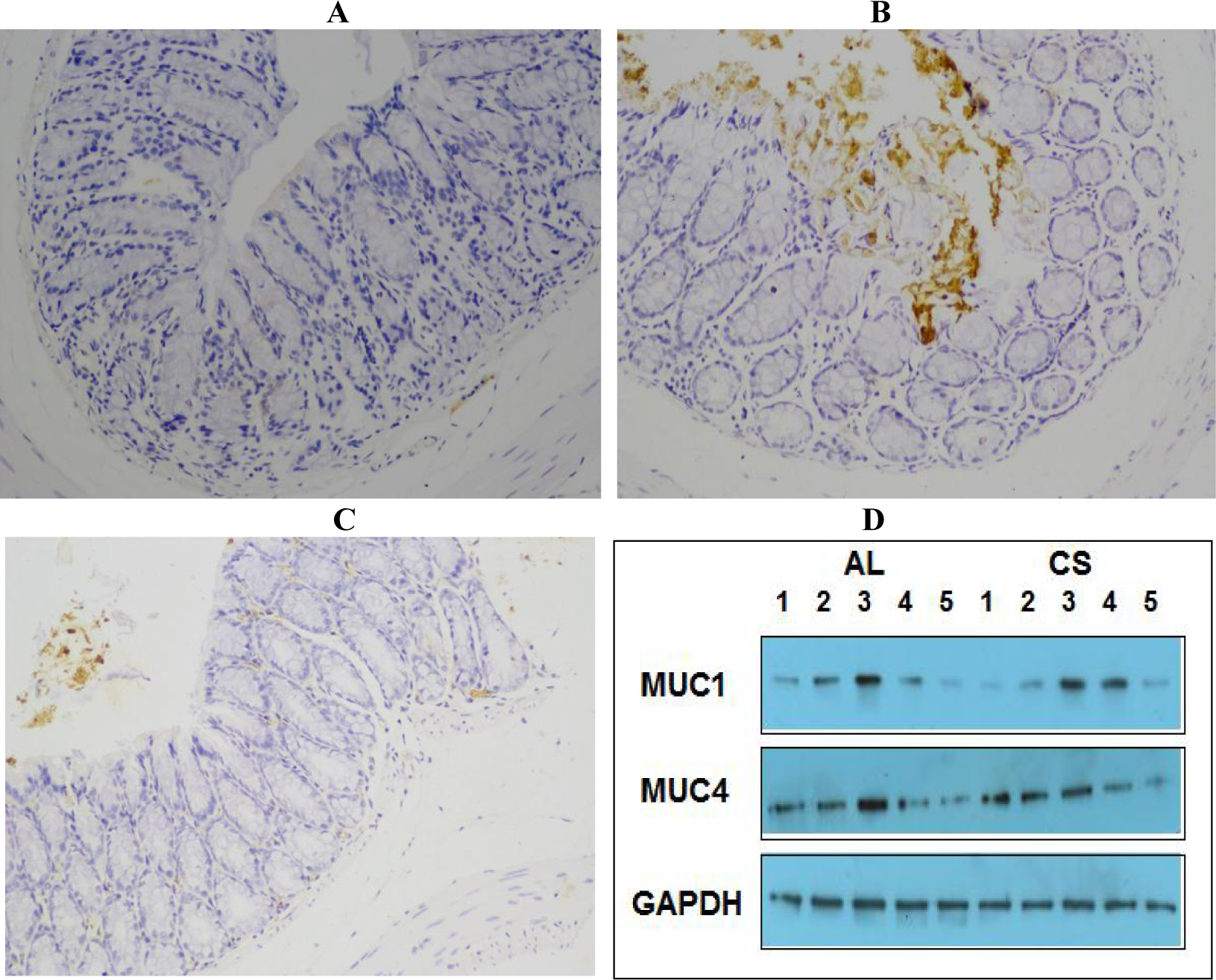
Mucus impairment and mucin expression in AL and CS mice. A. SPLI in the gut of AL2. B. SPLI in the gut of CS2. C. SPLI in the gut of CS3. D. MUC1/4 levels in AL and CS mice.

### CS induces tumor-like hyperplasia and tumor marker expression followed by a systemic pro-inflammatory response

During the large-scale evaluation of CS-induced pro-inflammatory responses, we unexpectedly observed a tumor-like hyperplasic object in the inner chest of a CS mouse (Fig. 6A). As compared with the circumjacent muscular tissues, the tumor marker CD1 shows a higher level in the neoplasm, whereas the tumor suppressor p21 exhibits a lower level in the neoplasm (Fig. 6B). To dissect the inflammatory profiles in this mouse carrying a neoplasm, we measured LPS and TNF-α levels in different tissues and the neoplasm *per se*. Intriguingly, LPS shows decline in hepatic and cardiac tissues, but elevation in adipose tissues (Fig. 6C). In similar, TNF-α is also declined in all tested tissues except for adipose tissues (Fig. 6D). Surprisingly, both LPS and TNF-α exhibit lower levels in the neoplasm. These results indicated that pro/anti-inflammation is differentially modulated in an organ-dependent manner, during which LPS and TNF-α in adipose tissues demonstrate a conversion from lower levels to higher ones, whereas those in other tissues also present a dynamic change from higher levels to lower ones.

**Fig. 6.**
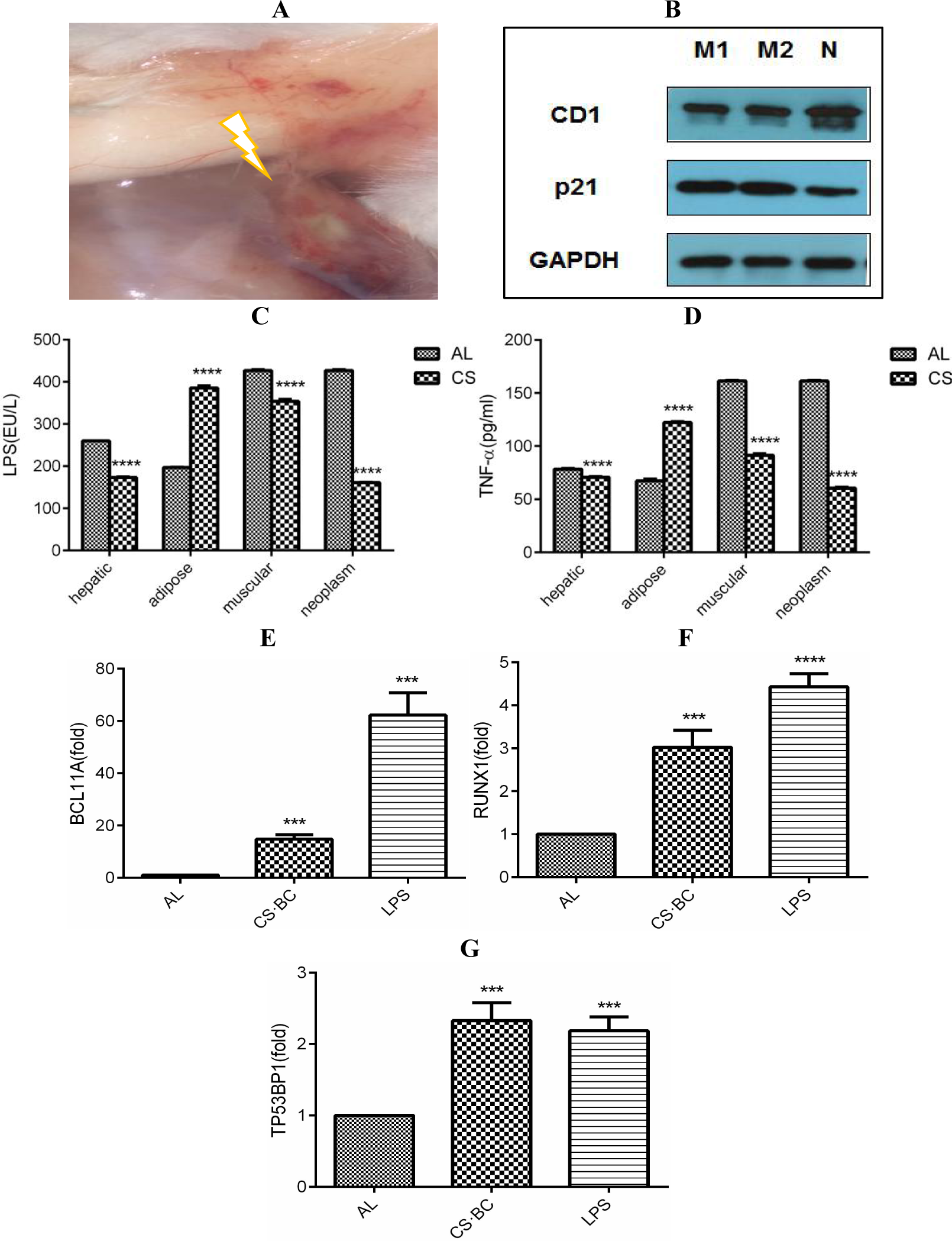
CS-induced tumor-like hyperplasia and tumor marker expression in AL, CS, CS-BC and LPS mice. A. Phenotypical morphology of tumor-like neoplasm. B. CD1 and p21 levels in the neoplasm, in which M1/M2 represents muscular tissues, and N represents a neoplasm sample. C. LPS levels in hepatic, adipose, muscular tissues and the neoplasm. D. TNF-α levels in hepatic, adipose, muscular tissues and the neoplasm. C-E. *RUNX1, TP53BP1* and *BCL11A* mRNA levels in mammary tissues. **** *p*<0.0001. *** *p*<0.001.

To further address an association of CS, SSB, LPS and tumorigenesis, we quantified the mRNA levels of three breast cancer-related markers, in the female mice fed with CS and intragastrically inoculated with *B. cereus*, or only intramuscularly injected with LPS. Consequently, the levels of *RUNX1, TP53BP1* and *BCL11A* mRNA as breast cancer markers are highly elevated in the mammary tissues of tested CS-BC and LPS mice (Fig. 6E-6G). These results indicated that SSB feeding or LPS administering can lead to the onset of mammalian tumorigenesis.

### CS regulates hypoxia, angiogenesis and mitochondrial biogenesis biomarkers

To explore whether inflammation would activate iNOS and HIF-1α to initiate the expression of downstream hypoxia-sensing genes, we measured the mRNA levels of iNOS, HIF-1α, VEGF and EPO in AL and CS mice. As depicted in Fig. 7A-7D, iNOS, HIF-1α, VEGF and EPO are upregulated in adipose tissues, but downregulated in the neoplasm of CS mice, suggesting adipose tissues in a pro-inflammatory status, but the neoplasm in an anti-inflammatory period. As an essential anti-inflammatory outcome, iNOS, HIF-1α, VEGF and EPO are downregulated globally in other tissues of CS mice.

**Fig. 7.**
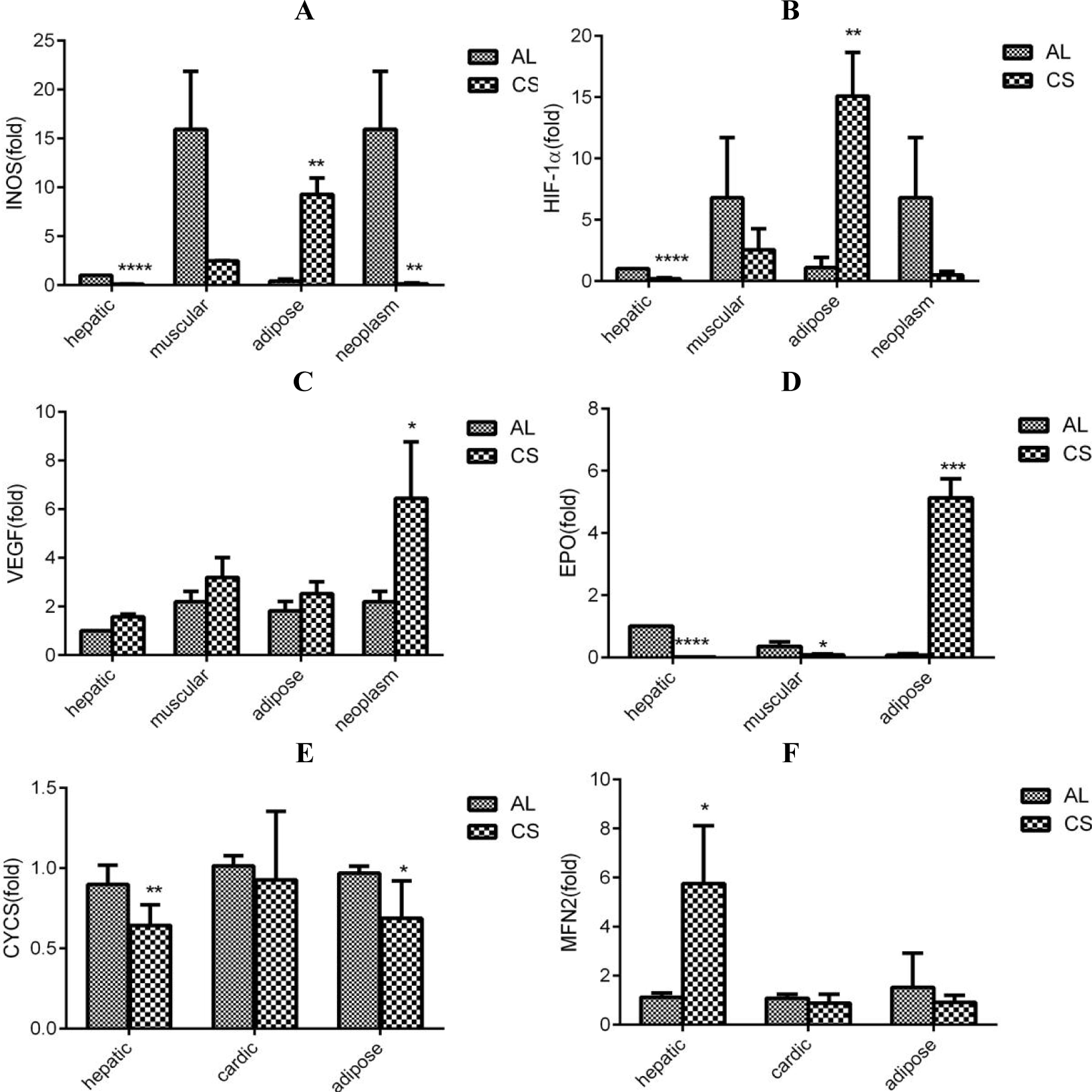
NO-driven hypoxic responses and mitochondrial biogenesis in different tissues and neoplasm of AL and CS mice. A. iNOS mRNA levels. B. HIF-1α mRNA levels. C. EPO mRNA levels. D. VEGF mRNA levels. E. CYCS mRNA levels. F. MFN2 mRNA levels. **** *p*<0.0001. *** *p*<0.001. ** *p*<0.01. * *p*<0.05.

On the other hand, CYCS and MFN2, as two mitochondrial biomarkers, are also differentially modulated in CS mice. As compared to AL mice, CSCS is downregulated in CS hepatic and adipose tissues, but unchanged in CS cardiac tissues (Fig. 7E). In contrast, MFN2 is upregulated in hepatic tissues, but unchanged in CS cardiac and adipose tissues (Fig. 7F). Those results derived from the adipose tissues suggested a compromised capacity of mitochondrial biogenesis that reflects a pro-inflammatory state in CS mice.

### CS-triggered inflammation induces fatty liver-related gene expression and fatty liver-like lipogenesis

To link CS-leaked LPS to hepatosteatosis and NALFD, we evaluated the expression profiles of 84 genes related to the pathogenesis to fatty liver in LPS-injected mice (**Additional file 3**). Consequently, genes for inflammatory responses and metabolic/signaling pathways were noticed to upregulation in the hepatic sample from a CS mouse (Table 4). For example, IL-1β is increased for 94 folds, IL-10 is increased for 50 folds, and IL-6 and TNF are increased for 12 and 10 folds. These results imply that hepatic inflammation might initiate a pathogenic process toward hepatosteatosis. Notably, carbohydrate metabolism responsible PDK4 and cholesterol metabolism/transport responsible ABCG-1 and CYP2E1 are increased for 21, 27 and 34 folds, respectively, hinting a trend towards NALFD.

**Table 4.** The hepatic expression levels of NAFLD-related genes with up/downregulation for more than two folds in AL and LPS mice

| Pathway/Metabolism | Gene | Description | Fold regulation of LPS to AL |
| --- | --- | --- | --- |
| Inflammatory response | *Cebpb* | CCAAT/enhancer binding protein (C/EBP), beta | 2.50 |
|  | *Ifng* | Interferon gamma | 5.52 |
|  | *Il10* | Interleukin 10 | 50.20 |
|  | *Il1b* | Interleukin 1 beta | 94.24 |
|  | *Il6* | Interleukin 6 | 12.54 |
|  | *Tnf* | Tumor necrosis factor | 10.24 |
| Carbohydrate metabolism | *G6pc* | Glucose-6-phosphatase, catalytic | 3.64 |
|  | *G6pdx* | Glucose-6-phosphate dehydrogenase X-linked | 3.27 |
|  | *Pck2* | Phosphoenolpyruvate carboxykinase 2 (mitochondrial) | 2.45 |
|  | *Pdk4* | Pyruvate dehydrogenase kinase, isoenzyme 4 | 21.27 |
|  | *Pklr* | Pyruvate kinase liver and red blood cell | -7.02 |
|  | *Rbp4* | Retinol binding protein 4, plasma | 2.65 |
| β-oxidation | *Acox1* | Acyl-coenzyme A oxidase 1, palmitoyl | 2.13 |
|  | *Cpt1a* | Carnitine palmitoyltransferase 1a, liver | 2.72 |
| Cholesterol metabolism/transport | *Abca1* | ATP-binding cassette, sub-family A (ABC1), member 1 | 6.48 |
|  | *Abcg1* | ATP-binding cassette, sub-family G (WHITE), member 1 | 27.10 |
|  | *Apoa1* | Apolipoprotein A-I | 2.83 |
|  | *Apob* | Apolipoprotein B | 6.97 |
|  | *ApoE* | Apolipoprotein E | 2.97 |
|  | *Cd36* | CD36 antigen | 2.37 |
|  | *Cyp2e1* | Cytochrome P450, family 2, subfamily e, polypeptide 1 | 34.38 |
|  | *Nr1h3* | Nuclear receptor subfamily 1, group H, member 3 | 3.39 |
| Insulin signaling pathway | *Igf1* | Insulin-like growth factor 1 | 3.19 |
|  | *Ptpn1* | Protein tyrosine phosphatase, non-receptor type 1 | 3.14 |
|  | *Socs3* | Suppressor of cytokine signaling 3 | 7.18 |
| Adipokine signaling pathway | *Cd36* | CD36 antigen | 2.37 |
|  | *LepR* | Leptin receptor | -2.60 |
|  | *Serpine1* | Serine (or cysteine) peptidase inhibitor, clade E, member 1 | 3.67 |
|  | *Slc2a1* | Solute carrier family 2, member 1 | 2.11 |
|  | *Stat3* | Signal transducer and activator of transcription 3 | 2.00 |

During evaluation of CS impacting on lipid accumulation (lipogenesis) or lipid degradation (lipolysis), we tried to observe whether oil deposits exist in the oil red-stained hepatic tissues. As compared with the hepatic sample from an AL mouse (Fig.8A), a large number of condensed oil droplets were seen in that from an CS mouse (Fig.8B). Accordingly, mitochondria that can degrade fatty acids via the Krebs cycle are extensively decreased in their numbers and sizes, which were distinguishable in muscular tissues (Fig. 8C and Fig. 8D) and adipose tissues (Fig. 8E and 8F) by less mitochondria and more oil droplets. These results suggested that systemic inflammation might have led to mitochondrial deficiency and metabolic dysfunction, by which impedes lipid catabolism and enhances lipid deposits.

**Fig. 8.**
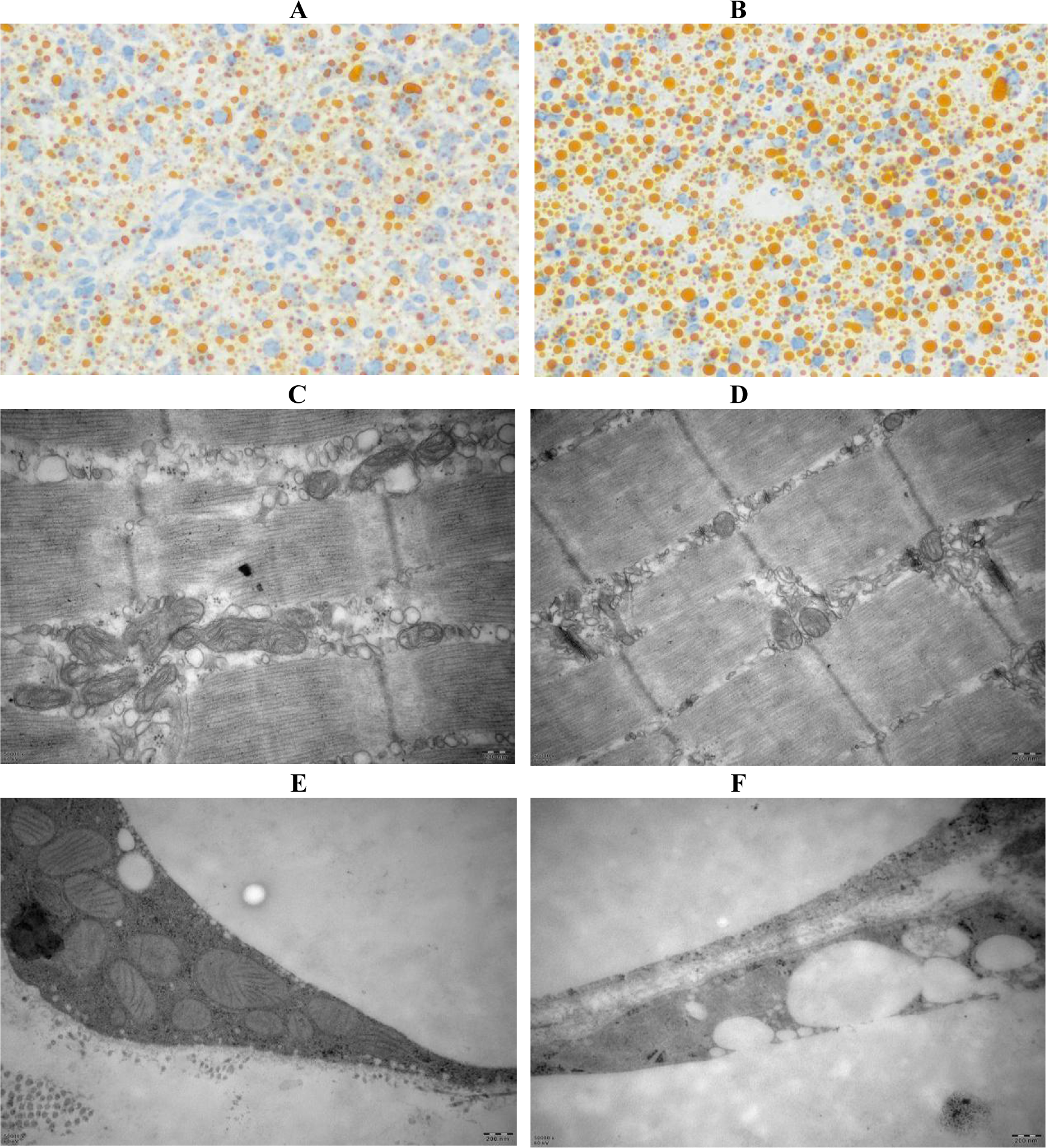
CS-induced fatty liver-like lipogenesis and NALFD-related gene upregulation in AL and CS mice. A. Oil red O stained hepatic tissues of an AL mouse (200×). B. Oil red O stained hepatic tissues of a CS mouse (200×). C. Mitochondrial morphology in muscular tissues of an AL mouse. D. Mitochondrial morphology in muscular tissues of a CS mouse (50000×). E. Mitochondrial morphology in adipose tissues of an AL mouse (50000×). F. Mitochondrial morphology in adipose tissues of a CS mouse (50000×).

## Discussion

Considering the emerging findings that CS induces sulfatase secretion from *B. thetaiotaomicron*, one of the bacterial species belonging to SSB, and thus nourishes *D. piger*, one of the bacterial species belonging to SRB [13], we hypothesized that CS-mediated gut opportunistic infection by SSB and SRB might be a potential trigger that reduces colon lining integrity, allows LPS leakage, induces chronic low-grade inflammation and initiates systemic pathogenesis towards multiple inflammatory diseases, including Alzheimer’s disease [24]. In the present study, we noticed for the first time that CS induces the early-phase and multi-systemic pathogenesis towards dementia, arthritis, tumor and fatty liver by modulating the corresponding hallmarks, i.e., Aβ upregulation for dementia, RF upregulation for arthritis, CD1 upregulation/p21 downregulation for tumor, and NAFLD-related gene overexpression for fatty liver. In particular, a synchronous decline of TH, Ach and ChAT levels was monitored in CS-fed mice, suggesting a hint initiating progressive cognitive deficits seen in LPS-challenged mice [25]. Moreover, arthritis-like erythematous and edematous paws, tumor-like neoplasia and fatty liver-like lipogenesis were also typically observed in CS-fed mice.

We also detected an overexpression of the transcription factor BCL11A that is overexpressed in the triplet negative breast cancer (TNBC) in the mammary tissues of CS-BC and LPS female mice. Recently, it was highlighted that exogenous BCL11A overexpression promotes tumor formation, whereas its knockdown suppresses tumorigenic proliferation, suggesting that BCL11A has a critically promoting role in TNBC [26]. As a newly identified prognostic indicator of TNBC that correlates with poor prognosis in TNBC patients [27], RUNX1 was also found upregulation in the mammary tissues of CS-BC and LPS female mice. Similarly, TP53BP1, an activator of the tumor suppressor p53 that is associated with TNBC [28], was also observed to have the raised mRNA levels in the mammary tissues of CS-BC and LPS female mice.

Our previous work indicated that live *Escherichia coli* feeding induces articular synovitis accompanying with the upregulation of iNOS and burst of NO [29]. Recently, *A. muciniphila* as a common member of SSB has been shown to allow mucus thinning, LPS leaking and epithelial proliferation [14]. As evidence supporting LPS-mediated neurodegenerative disease, a species of gut bacteria was found to be correlated to the clinical phenotype of Parkinson’s disease [11]. It was also revealed that LPS and K99 derived from Gram negative bacteria exist in the amyloid plaques of dementia patients’ brain [30]. Furthermore, it was verified a pathogenic mechanism underlying a specific gut microbiota induces neuroinflammation and causes cognitive degeneration in patients with Parkinson’s disease [31].

In this experimental observation, we unambiguously noted a coherent changes of sulfatase and mucin expression with leukocyte activation. When sulfatase is overexpressed, SLPI is upregulated, and MUC1/4 is accordingly overproduced. In other words, mucus repairs are enhanced after colon lesions. By nourishing SSB, CS was found to elevate serum LPS levels and upregulate TNF-α in multiple tissues, implying gut leakage and LPS entry into the blood circulation. As similar as bacteremia leading to sepsis and septic shock, LPS entering into the bloodstream can trigger a potent immune response in a titer-dependent manner. While high-titer LPS resembling acute bacterial infection induces neutralizing antibodies for elimination of LPS, low-titer LPS resembling chronic bacterial infection slightly elevates the levels of pro-inflammatory cytokines for two to three folds, leading to low-grade inflammation and initiating diabetes mellitus and fatty liver diseases [23].

Interestingly, we observed that LPS and TNF-α levels in adipose tissues are different from those in other tissues. Briefly, LPS and TNF-α levels in adipose tissues are declined at first, but elevated later. In contrast, LPS and TNF-α levels in other tissues are first elevated, but then declined. This intriguing phenomenon seemed to be explained by a finding that SRB1 binds to LPS and incorporates it into chymicrons, which are later enhanced transcytosis over the endothelial barrier and endocytosis of adipocytes [32]. From this clue, we deciphered that adipose tissues should uptake high-level LPS and enhance antibody induction and LPS eradication. Due to the interaction of SRB1 with LPS, the concentration of LPS must eventually be higher than the titer of anti-LPS, which should allow LPS accumulation in adipose tissues to elevate TNF-α levels. Without a specific role of LPS-SRBI in other tissues, the concentration of LPS could not be higher than the titer of anti-LPS, so LPS and TNF-α should be converted from higher levels to lower ones.

Logically, if serological tests would be performed, the inflammatory indicators should be lower (in an anti-inflammatory phase). In contrast, if tests would be conducted using a tissue sample, the inflammatory indicators should be higher (in a pro-inflammatory stage). Accordingly, we found in the present study that LPS upregulates the NALFD-related genes in a large scale. However, our recent work indicated that most of the NALFD-related genes are downregulated in CS mice as compared with AL mice [33]. The latter result addressed that CS might not raise the hepatic LPS level to upregulate the NALFD-related genes, otherwise it should have reduced the hepatic LPS level to downregulate the NALFD-related genes. Indeed, this explanation can be supported by the low hepatic LPS level in CS mice tested in this study, suggesting that the residual LPS might have been eventually eradicated by the neutralizing anti-LPS antibody. Nevertheless, some other NALFD-related genes were still observed to be upregulated in a different manner and an unknown mechanism.

It has been documented that CS shows differential results in dealing with OA, i.e., some reports demonstrating effective [15,16], but others addressing ineffective [18,19]. For this discrepancy, it might be simply deciphered by an individual gut microbiota specificity. CS should exert anti-inflammatory effects and ameliorate OA when SSB are absent or rare in the gut, otherwise it should exert pro-inflammatory effects and aggravate OA. In the present study, we noticed a synovitis-like phenotype in CS.RA with the overgrown SSB (*B.cereus*), but not in CS.HL without the overgrown SSB. However, it remains inclusive why CS.NM with the overgrown SSB (*A.muciniphila*) does not show synovitis. In our recent finding, *A.muciniphila* was observed to induce mucin expression and reinforce mucus integrity, thereby compromising LPS-mediated pro-inflammatory stimuli [33]. Accumulating evidence supports that *A. muciniphila* has therapeutic values in metabolic diseases, including an inverse correlation of *A. muciniphila* with obesity and diabetes in mice and humans [34, 35].

On the other hand, a pro/anti-inflammatory role of CS on OA might be also explained by a time-dependent condition, in which CS is effective against OA once LPS is eradicated by anti-LPS in a later stage, otherwise OA should be deleterious because LPS levels are higher in an early period. In this study, we found that most tissues in CS mice exhibit the pro-inflammatory responses at first, but then exert an anti-inflammatory effect later. Accordingly, LPS and TNF-α levels are higher in an early phase of CS feeding, but both levels are declined in the late period. These results indicated that CS may eventually show an anti-inflammatory effect by raising LPS levels to induce anti-LPS generation in almost all tissues except for adipose tissues. In our previous work, we surprisingly found that a relative higher dose of LPS (1.2 mg/kg) leads to a peritoneally anti-inflammatory state, whereas a relative lower dose of LPS (0.25 mg/kg) causes a viscerally pro-inflammatory state, implying anti-LPS neutralizing LPS in peritoneal organs but not in visceral organs [23].

LPS can bind to TLR4 on cell surface to activate NF-κB for upregulation of pro-inflammatory cytokines including TNF-α and IL-1β [36]. Actually, we detected the synchronous upregulation of LPS, TNF-α and TNFR1. After depletion of LPS, iNOS-derived high-level NO should be blocked, and eNOS-derived NO would be rehearsed, thereby activating adenosine monophosphate-activated protein kinase (AMPK) and initiating mitochondrial biogenesis [37]. Alternatively, AMPK can in turn activate eNOS by the signaling cascade AMPK→ Rac1→Akt→eNOS [38]. Furthermore, it has been raveled that AMPK can catalyze the phosphorylation of JAK2 and inhibit its pro-inflammatory activity via JAK2-STAT3 signaling [39]. The extracellular infection by pathogenic microorganisms can activate NADP oxidase to trigger the burst of reactive oxygen species (ROS), which upregulates the pro-inflammatory cytokine expression and augment the pro-inflammatory response [40]. It was accepted that TNF-α, IL-1β and other pro-inflammatory cytokines can upregulate iNOS for NO production, and that NO as a hypoxic inducer can activate HIF-1α [41,42].

In our previous work [29], we showed that the NO donor sodium nitroprusside can activate HIF-1α and upregulate hypoxia-responsive genes such as VEGF. On the other hand, EPO, a glycoprotein hormone indispensable for erythropoiesis, was validated to reduce blood glucose and body mass in mice [43]. Additionally, EPO was thought to have other biological activities that extend to nonerythroid tissues, such as inhibition of adipose inflammation, normalization of insulin sensitivity, and reduction of glucose intolerance [44]. In this investigation, we observed the elevation of iNOS, HIF-1α, EPO and VEGF levels in adipose tissues of CS mice, suggesting a hypoxic milieu leading to mitochondrial dysfunction and deficiency among other abnormal outcomes. Accordingly, we detected the downregulation of MFN2 and CSCS, two common mitochondrial biomarkers [45], in the most tested tissues of CS mice, implying compromised mitochondrial biogenesis and mitigated lipid degradation. Indeed, we noticed that the numbers and sizes of mitochondria are dramatically decreased in muscular and adipose tissues of CS mice. We also found that a large number of condensed oil droplets are accumulated in hepatic tissues by CS feeding, perhaps due to without sufficient mitochondria for fatty acid degradation.

## Conclusions

The animal-based dietary sulfate CS would initiate the onset of multi-systemic inflammatory pathogenesis by nourishing the mucin-degrading microbiota and compromising the mucus-protecting function. CS might be beneficial or harmful to OA depending on the absence or presence of SSB and SRB. Upon interaction with SSB and SRB, CS induces multi-systemic pathogenesis towards arthritis, dementia, tumor and fatty liver although they maintain in a mild state and during an early stage.

## List of abbreviations

Aβ: amyloid β peptide
ACh: acetylcholine
AL: *ad libitum* chow
AMPK: adenine monophosphate-activated protein kinase
CCL2: chemokine (C-C motif) ligand 2
CD1: cyclin D1
ChAT: choline acetyltransferase
CS: chondroitin sulfate
CSCS: cytochrome *c*, somatic
ELISA: enzyme-linked immunosorbent assay
EPO: erythropoietin
GAPDH: glyceraldehyde-3-phosphate dehydrogenase
HE: haematoxylin-eosin
HIF-1α: hypoxia induced factor 1α
LPS: lipopolysaccharide
IL-1: interleukin 1
iNOS: inducible nitric oxide synthase
MFN2: mitofusin 2
MUC: mucin
NALFD: non-alcoholic fatty liver disease
NO: nitric oxide
PCR: polymerase chain reaction
PS-1: presenilin 1
qPCR: quantitative real-time PCR
RF: rheumatoid factor
TH: tyrosine hydroxylase
TLR4: toll-like receptor 4
TNBC: triplet negative breast cancer
TNF-α: tumor necrosis factor α
TNFR1: TNF-α receptor 1
SLPI: secretory leukocyte protease inhibitor
SRB: sulfate-reducing bacteria
SSB: sulfatase-secreting bacteria
VEGF: vascular epithelial growth factor
WB: Western blotting.

## Declarations

### Ethics approval and consent to participate

All subjects signed written consent to participate in this study. This study was approved by The Animal Care Welfare Committee of Guangzhou University of Chinese Medicine (No. SPF-2015009).

### Consent for publication

Not applicable.

### Availability of data and materials

The datasets supporting the conclusions of this article are included within the article (**Additional file 1 includes primer sequence data, Additional file 2 includes gut microbiota metagenomic data, and Additional file 3 includes fatty liver-related gene microarray data**).

### Funding

This work was supported by the National Natural Science Foundation of China (No. 81273620 to

Qing-Ping Zeng, No. 81673861 to Chang-Qing Li, and No. 81273817 and No. 81473740 to Qi Wang), and Guangdong Science and Technology Plan Project (No. 20150404042 to Qin Xu).

## Acknowledgments

We thank our colleagues in the Tropical Medicine Institute and Clinical Pharmacology Institute, Guangzhou University of Chinese Medicine, China. We also thank Kangchen Biotechnology Co, Shanghai, China for performance of RT-PCR array experiments, and Novogene, Beijing, China for conduction of gut microbiota metagenomic analysis.

## Authors contributions

QPZ, QW and QX designed the study. TL, YPC, and LLT carried out the experiments. XAH performed bioinformatics analysis. CQL, QW and QX participated in the interpretation of results. QPZ wrote the manuscript with input from other authors. All authors read and approved the final manuscript.

